# Dishevelled drives Wnt-stimulated disassembly of primary cilia through a unique PDZ-mediated binding mode with Daple and Girdin

**DOI:** 10.1101/2025.09.26.678488

**Authors:** Miha Renko, Gonzalo J. Beitia, Trevor J. Rutherford, Mariann Bienz, Melissa V. Gammons

## Abstract

Dishevelled is a pivotal cytoplasmic hub protein that transmits Wnt signals to various cytoplasmic effectors to specify cell fates and behaviors during animal development. The molecular mechanisms by which Dishevelled directs Wnt outputs towards β-catenin or other non-canonical effectors remain unclear. Its PDZ domain is dispensable for signaling to β-catenin but essential for multiple non-canonical Wnt responses in Drosophila and vertebrate systems. None of its functionally relevant binding partners are known even though a broad range of PDZ-binding ligands have been identified. Here, we combined proximity labeling with structural and biophysical analysis to discover that Daple and its Girdin-L paralog bear unique extended C-terminal PDZ-binding motifs that bind to the PDZ domain of the main human Dishevelled paralog DVL2 with exceptionally high affinity. Assays in HEK293T cells revealed that deletions of these motifs or their cognate PDZ domain of DVL2 resulted in elongated primary cilia and rendered these cilia unresponsive to Wnt5a-stimulated disassembly following serum starvation. We conclude that an unprecedented molecular interaction between Dishevelled and Daple or Girdin-L underpins the disassembly of these ciliary organelles with universal links to signaling and cell cycle progression.

**One-sentence summary:** The PDZ domain of Dishevelled engages in unique interactions with the C-termini of Daple or Girdin-L to mediate the disassembly of primary cilia in response to Wnt5a.

## INTRODUCTION

Wnt signaling pathways operate in all animals and humans to control cell fates and tissue patterning (*1–3*). Canonical Wnt ligands signal through β-catenin, typically to promote cell proliferation and tissue homeostasis (*4*), whereas non-canonical Wnt signaling branches tend to coordinate behaviors of groups of cells such as their planar polarization or migration(*5*).

All Wnt signals pass through the cytoplasmic hub protein Dishevelled where signals diverge into β-catenin-dependent or non-canonical effector branches (*6*). How Dishevelled selects different effectors in response to different Wnt ligands remains unclear, although the Wnt transmembrane co-receptors are thought to be pivotal for this choice.

To transduce Wnt signals, Dishevelled must bind to the intracellular face of a seven-transmembrane Frizzled receptor through its DEP (Dishevelled, Egl-10 and Pleckstrin) domain(*7–9*). This interaction is triggered by Wnt binding to the extracellular domain of Frizzled and its coupling to one of several Wnt co-receptors (*3*). The formation of a Wnt receptor complex involves Wnt-induced dimerization or tetramerization of Frizzled (*10*, *11*), which enables Dishevelled to engage in dynamic head-to-tail polymerization through its DIX (Dishevelled and Axin) domain (*12*) to assemble signalosomes with a high binding avidity for various low-affinity effector proteins (*1*, *13*).

Dishevelled also contains a PDZ (**P**ost-synaptic density protein-95, **D**iscs large tumor suppressor, **Z**onula occludens-1) domain which is dispensable for β-catenin-dependent Wnt signal transduction in human cells as shown in physiological complementation assays based on stable re-expression of wild-type (WT) and mutant DVL2 at near-endogenous levels (*14*). This is also true in *Drosophila* where genomic excision of the PDZ domain of *dishevelled* (*dsh*) revealed that Armadillo-dependent Wingless signaling was intact while uncovering a range of tissue-specific non-canonical Wnt signaling defects (*15*). Indeed, the Dishevelled PDZ domain is essential for non-canonical Wnt5a-dependent activation of the downstream effectors, Rac and Rho GTPases, to drive cell migration by binding to Daple and Daam1, respectively (*16*, *17*). Together, these results indicate that the PDZ domain of Dishevelled represents a key interaction module for the transduction of non-canonical Wnt signals.

PDZ domains are small globular protein-protein interaction modules (*18*) that recognize PBMs (**P**DZ-**b**inding **m**otifs) typically found at the C-terminus of their ligands (*19*) through a peptide binding groove formed by its β2 strand, its α2 helix and its highly conserved carboxylate binding loop (*20*, *21*). PBMs typically consist of three C-terminal residues that were initially categorized into three main classes, namely Ser/Thr-X-Φ (class I), Φ-X-Φ (class II) and Glu/Asp-X-Φ (class III) where Φ is hydrophobic and X is any amino acid (*22*, *23*). However, subsequent studies have shown that some PDZ domains can accommodate longer PBMs (*24*, *25*). Indeed, the PDZ domain of DVL2 (PDZ_DVL2_) has a highly adaptable and extended ligand-binding cleft owing to its elongated β2 strand and α2 helix, enabling it to accommodate numerous diverse ligands of up to seven residues (*26*) including a subset of class I and class II PBMs (*14*, *15*, *27*). In support of this, a comprehensive analysis aimed at establishing specificity maps for 72 recombinant human and worm PDZ domains revealed that Dishevelled PDZ domains preferentially bind to extended class 1-like PBMs (Lys-Ω-Φ-Gly-Trp-Phe where Ω is aromatic) (*24*). Taken together, these findings illustrate the versatility and selectivity of Dishevelled PDZ domains in binding to and discriminating between multiple PBMs owing to their extended binding cleft.

Although biophysical assays are powerful tools for identifying ligands of individual PDZ domains, it is a considerable challenge to identify those that are biologically relevant. For example, a previous biophysical study identified three components of the distal polarity core complex (Dishevelled, Frizzled and Starry Night) as ligands of PDZ_DVL2_, but the combined genetic deletion of the genes encoding these components in *Drosophila* failed to recapitulate the non-canonical Wnt signaling defects of flies lacking the PDZ domain of its single Dishevelled ortholog (*15*). Therefore, it remains unclear which of the reported ligands of PDZ_DVL2_ mediate its essential roles in transducing non-canonical Wnt signals.

Here, we used BioID proximity labeling to identify Girdin and its Daple paralog as the top interactors of PDZ_DVL2_ in human cells. Structural and biophysical analyses revealed a unique binding mode where all eight residues of the Daple PBM interact with this domain, consistent with its unusually high binding affinity for PDZ_DVL2_. By truncating the PBMs from endogenous Daple and a Girdin isoform called Girdin-L (*28*) with CRISPR/Cas9 gene editing, we discovered defects in the disassembly of primary cilia that were also evident in cells lacking all three human Dishevelled paralogs (DVL1-3). These ciliary defects were rescued by re-expression of WT DVL2 at physiological levels but not by DVL2 lacking its PDZ domain. Furthermore, we found that the Wnt5a-stimulated disassembly of primary cilia depended on the PBMs of Daple and Girdin-L and on the structural integrity of their cognate PDZ_DVL2_. We thus uncovered an essential role of this domain in the disassembly of primary cilia that is mediated by PDZ_DVL2_ and its unique interaction with Daple and Girdin-L.

## RESULTS

### Daple and Girdin-L engage in strong interactions with PDZ_DVL2_ through their PBMs

To identify the interactome of PDZ_DVL2_ in the human epithelial kidney cell line HEK293 (Flp-In T-Rex 293), we used a proximity-labeling approach called BioID, tagging WT DVL2 and DVL2 lacking its PDZ domain (ΔPDZ) with BirA* (a promiscuous version of the biotin ligase BirA) (*29*). We identified 30 hits whose labeling consistently depended on the presence of PDZ_DVL2_ across three independent experiments regardless of exogenous Wnt stimulation, including VANGL1 (*30*, *31*) and Daple (*16*, *32*, *33*) that were previously implicated in Wnt signaling (**Fig. 1A, B & Table S1**). In addition, we identified several proteins localized to centrosomes or primary cilia (e.g. CEP192, ALMS1, OCRL), including Girdin (*34*), a close relative of Daple that localizes to the centrosomes and basal bodies (*35*, *36*) which nucleate the growth of primary cilia. Of note, many of these hits reflect vicinal proteins that do not interact directly with PDZ_DVL2_, which may explain why more than half of them lack a recognizable C-terminal PBM (**Fig. 1B & Table S1**).

**Fig. 1.**
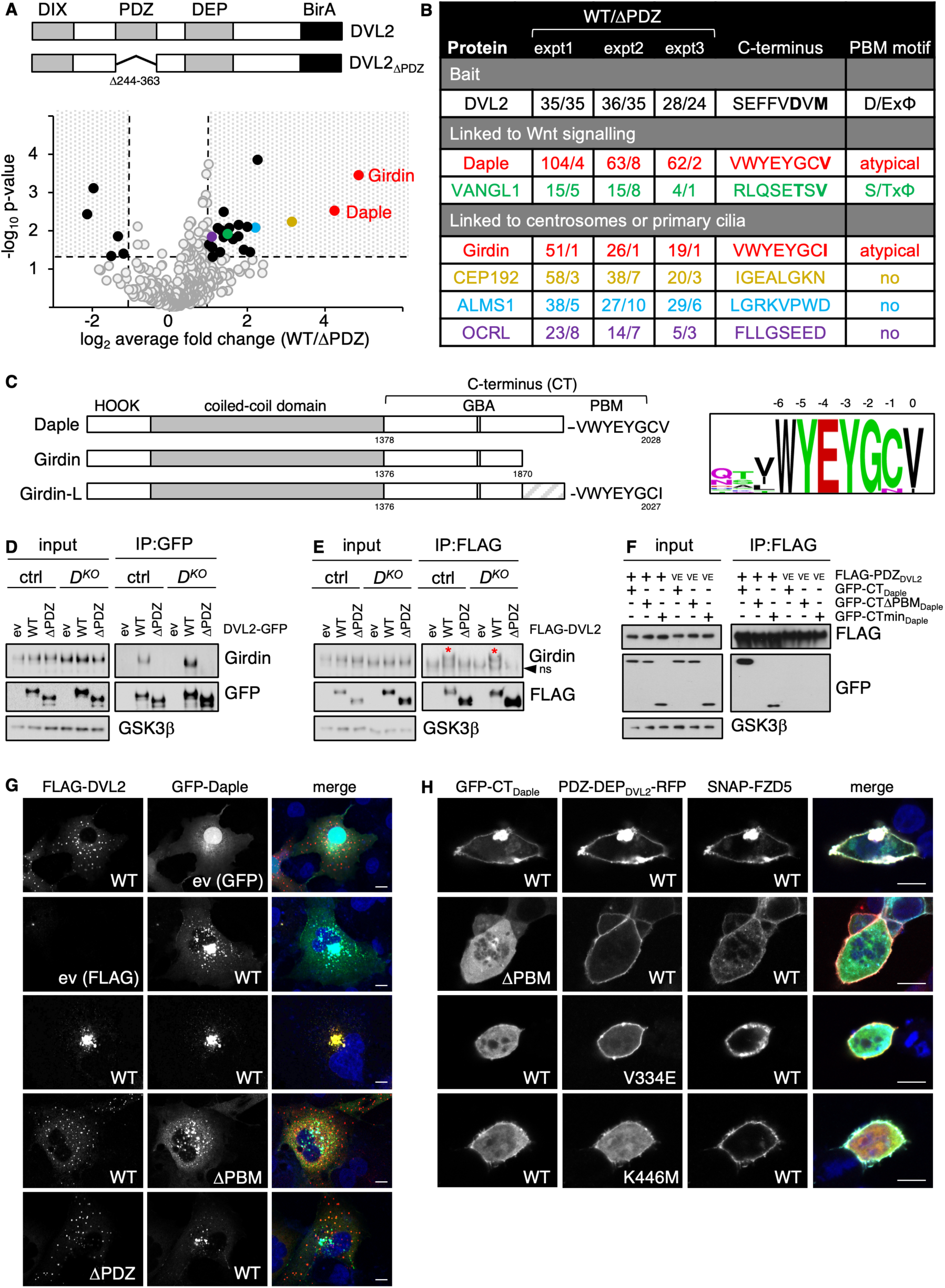
Girdin and Daple bind to the PDZ domain of DVL2. (**A**) Volcano plot of PDZ_DVL2_-sensitive BioID hits; n=3 biological repeats; above, cartoon of DVL2-BirA constructs. (**B**) List of selected PDZ_DVL2_-sensitive hits (see also **Table S1**); numbers represent unweighted exclusive unique spectral counts (>95% probability). (**C**) *Left*, domain architecture of Daple, Girdin and Girdin-L with conserved motifs (GBA, G protein-binding and -activating motif; PBM, PDZ-binding motif). *Right*, JACKHMMER plot of the C-terminal sequences of mammalian Daple and Girdin orthologs. (**D, E**) Western blots of immunoprecipitates (IPs) from HEK293T cells (ctrl) or Daple-knockout (*D^KO^*) cells transiently transfected with wild-type (WT) or mutant DVL2 tagged with N-terminal FLAG or C-terminal GFP (as indicated on blot), probed with antibodies as indicated on the right; representative of *n=3* biological repeats. (**F**) Western blots of coIPs of WT or V334E (VE) mutant FLAG-PDZ_DVL2_ after co-expression with WT or GFP-CT_Daple_ or GFP-CTmin_Daple_ in transiently transfected HEK293T cells, probed with antibodies as indicated on the right; representative of n=3 biological repeats. (**G**) Representative confocal images selected from >100 HeLa cells co-expressing WT or mutant FLAG-DVL2 and GFP-Daple, as indicated in panels. (**H**) Representative confocal images selected from >100 transfected HEK293T cells, co-expressing SNAP-FZD5 with WT or mutant GFP-DapleC and WT or mutant PDZ-DEP_DVL2_-RFP, as indicated in panels. Scale bars, 10 μm.

Daple and Girdin can be considered paralogs as they both contain an N-terminal HOOK and a central coiled-coil domain, including a leucine zipper that mediates self-association, and an unstructured C-terminal tail (CT) bearing a G protein α-binding and -activating motif (GBA) and an atypical class I-like PBM at their C-termini(*16*, *32*) (**Fig. 1C**). In mammalian cells, Daple binds to PDZ domains of different DVL paralogs through its PBM to promote non-canonical Rac-dependent Wnt responses while antagonizing canonical Wnt signaling (*16*, *32*). However, its CT diverges considerably from that of Girdin, and the latter was initially thought to lack a C-terminal PBM (*16*, *32*). However, an extended splice variant retaining introns 31 and 32 was since discovered (*28*) whose carboxy terminus closely resembles that of Daple. Indeed, the extreme C-termini of Daple and Girdin-L are almost identical, differing only by the last residue (Val in Daple, Ile in Girdin-L; **Fig. 1C & Fig. S1**). They are also deeply conserved throughout evolution and unique to these two proteins, which suggests their functional relevance. Although we do not know which Girdin isoform was identified as one of the top PDZ-sensitive hits in our BioID screen, it seems likely that this hit corresponded to the Girdin-L isoform which may bind directly to PDZ_DVL2_ through its atypical PBM.

To test this, we used co-immunoprecipitation (coIP) assays in HEK293T cells transiently transfected with GFP-or FLAG-tagged DVL2, to monitor its association with endogenous Girdin. Indeed, Girdin readily coIPed with WT DVL2, but not with ΔPDZ, regardless of the tag (**Fig. 1D, E**). We also performed coIPs in Daple knockout (*D^KO^*) cells generated by CRISPR/Cas9 gene editing, to rule out that Girdin associated with Dishevelled through endogenous Daple. The results were essentially the same: endogenous Girdin coIPed with WT DVL2 but not with ΔPDZ (**Fig. 1D, E**). Therefore, Girdin associated with DVL2 in a PDZ-dependent manner, like Daple (*16*, *32*).

Next, we confirmed that FLAG-DVL2 associates with Daple through its C-terminal PBM in transiently transfected HEK293T cells (*16*, *37*, *38*) (**Fig. S2A**). We also tested whether the minimal PDZ domain of DVL2 (PDZ_DVL2_, amino acids 265-361) is sufficient to associate with the CT of Daple (CT_Daple_, amino acids 1378-2028) or a minimal C-terminal polypeptide bearing its PBM (CTmin_Daple_, amino acids 1925-2028). This was the case (**Fig. 1F**): GFP-CT_Daple_ and GFP-CTmin_Daple_ coIPed with WT PDZ_DVL2_ but not with a mutant domain bearing a V334E substitution in its binding cleft that blocks binding to PDZ ligands (*14*).

Furthermore, FLAG-PDZ_DVL2_ failed to coIP with the minimal Daple CT that lacked its PBM (GFP-CTΔPBM_Daple_; **Fig. 1F**). These results from our coIP assays indicated a strong interaction between the two proteins mediated entirely by the C-terminal PBM of Daple and its cognate PDZ domain in DVL2.

### Recruitment of Daple to Frizzled is mediated by Dishevelled

Previously, it was thought that Daple binds directly to the C-terminal tails of several FZD paralogs, with a strong preference for FZD7 (*33*). These authors also proposed that an internal PBM of FZD7 competes with the Daple PBM in their binding to the PDZ domain of DVL1 (*33*). However, it has since been established beyond doubt that DVL binds through its DEP rather than its PDZ domain to the cytoplasmic face of FZD receptors (*7*, *8*, *39*).

Furthermore, although PDZ_DVL2_ can bind to the class I PBM found at the C-termini of most FZD paralogs (*15*), this is a weak interaction that does not suffice to mediate recruitment of DVL to the FZD receptor complex (*8*, *39*). Of note, PDZ_DVL2_ is dispensable for canonical Wnt signaling (*14*, *15*) which argues strongly against its function in Frizzled binding.

To demonstrate the critical role of the DEP domain of DVL2 in the recruitment of Daple to FZD receptors, we used a well-characterized DEP-dependent recruitment assay based on based on immunofluorescence assays in HeLa cells transiently co-transfected with SNAP-tagged Frizzled (*7*, *8*, *39*). If expressed on their own, murine GFP-Daple accumulated at the microtubule organising centre (MTOC) and in perinuclear regions, as previously observed (*40*), while FLAG-DVL2 formed distinct cytoplasmic puncta (*41*), but the two proteins co-localized if WT albeit not mutant proteins (lacking their PDZ domain or their C-terminal eight amino acids, ΔPDZ or ΔPBM, respectively) were co-expressed (**Fig. 1G**), confirming that their association depends on a PDZ-PBM interaction. Similarly, GFP-CT_Daple_ was cytoplasmic and diffuse when expressed by itself in HEK293T cells, but was efficiently recruited into FLAG-DVL2 puncta in a PBM-dependent manner (**Fig. S2B**). However, the recruitment of GFP-CT_Daple_ to membrane-localized SNAP-FZD5 strictly depended on co-recruitment of an RFP-tagged DVL2 fragment spanning its PDZ and DEP domains (PDZ-DEP_DVL2_-RFP) which is capable of binding to both FZD5 and CT_Daple_ (**Fig. 1H**). Membrane co-recruitment of GFP-CT_Daple_ was abolished by deletion of its PBM, or by mutations in the PDZ cleft (called V334E), or in the Frizzled-binding DEP ‘finger tip’ of PDZ-DEP_DVL2_-RFP (called K446M) that blocks its binding to Frizzled (*7*, *8*, *39*) (**Fig. 1H**). These results confirmed that the DEP and PDZ domains of DVL2 can engage in simultaneous independent interactions with their cognate ligands. They also argue against the previous claim (*33*) of a direct physical interaction between Frizzled and Daple.

### The PBMs of Daple and Girdin-L undergo extensive interactions with PDZ_DVL2_

The C-terminal sequences of Daple and Girdin-L do not conform to any previously identified PBM consensus motif (*24*, *25*). However, the C-terminus of Daple contains a PBM-like motif that was thought to mediate its binding to PDZ_DVL2_ in cells (*16*), which we have confirmed (**Fig. S2A, B**), but its direct binding to PDZ_DVL2_ has never been shown. We therefore tested peptides spanning their C-terminal eight residues (PBM_Daple_, PBM_Girdin_) for binding to PDZ_DVL2_ using nuclear magnetic resonance (NMR) spectroscopy (*14*, *15*). We purified bacterially expressed ^15^N-labelled PDZ_DVL2_ and acquired ^1^H-^15^N correlation spectra, alone or after incubation with purified PBM_Daple_ and PBM_Girdin_ tagged with Lipoyl (Lip, to increase solubility).

Both peptides caused a large number of chemical shift perturbations (CSPs) or line broadenings (spectral perturbations referred to as ‘bleaching’ below) relative to unliganded PDZ_DVL2_ (**Fig. 2A, B**). Indeed, a greater number of CSPs and bleaching events were observed compared to other peptides known to bind PDZ_DVL2_, such as Lip-PBM_FZD4_ (*15*) (**Fig. 2C**), Lip-PBM_VANGL1_ (**Fig. S3A**) or other previously tested ligands all of which comprise canonical class I-III PBMs(*14*, *15*). Next, we incubated Lip-PBM_Daple_ with the binding-defective V334E cleft mutant (*14*), which reduced the numbers of CSPs and bleaching events but did not abolish them (**Fig. S3B**). This was different from canonical PBMs previously tested in the same NMR assay whose interactions with PDZ_DVL2_ was abolished by V334E (*14*, *15*), suggesting that the Daple PBM engages in additional interactions outside the main hydrophobic anchoring pocket of the PBM cleft.

**Fig. 2.**
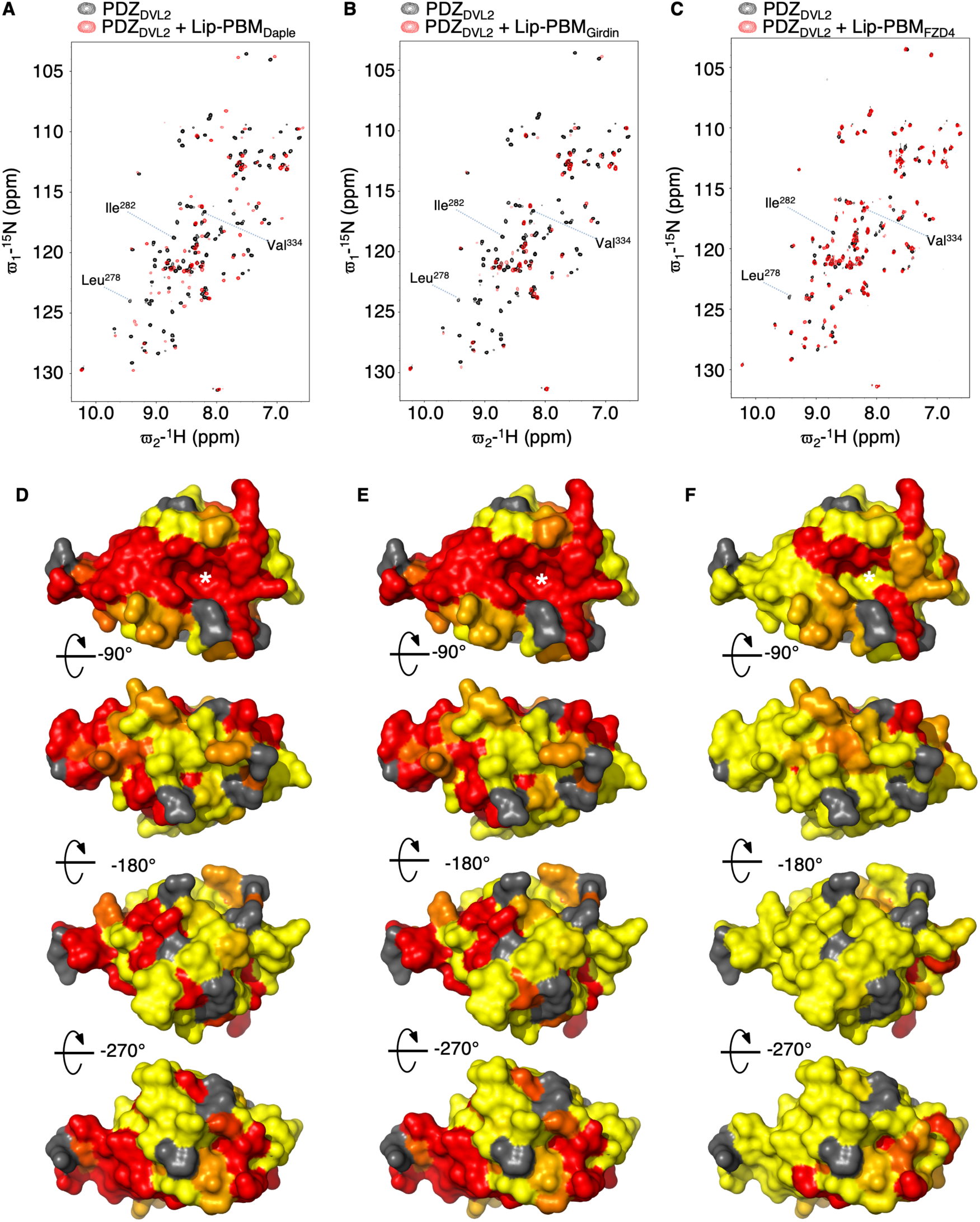
The PBMs from Daple and Girdin-L engage in extensive interactions with PDZ_DVL2_. (**A-C**) Overlay of HSQC spectra of 100 μM ^15^N-labelled PDZ_DVL2_ alone (*black*) or incubated with (**A, B**) 150 μM or (**C**) 300 μM Lip-tagged PBM peptides (*red*), as indicated above panels. (**D-F**) Corresponding heat-maps of line broadening [*red*, >mean+1 SD (1σ) fractional reduction of peak height] and chemical shift perturbations (*orange*, >mean+1σ |Δδ1H|+|Δδ15N/5|) projected onto the crystal structure of PDZ_DVL2_ (PDB: 2REY) in surface representation; *white asterisks*, hydrophobic pockets of PBM-binding cleft.

Using previous resonance assignments of ^15^N-^13^C double-labeled PDZ_DVL2_ (*14*), we were able to generate ‘heat-maps’ where each CSP or bleaching event indicates that the corresponding PDZ_DVL2_ residue either undergoes an interaction with our Lip-tagged peptides or changes its local structural environment. Indeed, a heat-map of Lip-PBM_Daple_ and Lip-PBM_Girdin_ projected onto the crystal structure of PDZ_DVL2_ (PDB: 2REY) confirmed that, in both cases, several of the CSPs induced by these ligands involve residues that are located outside its PBM-binding anchoring pocket (**Fig. 2D, E**). This was neither the case for Lip-PBM_FZD4_ (**Fig. 2F**) nor for previously tested PBMs (*14*, *15*) whose CSPs were essentially confined to the residues that form this pocket. This is consistent with the notion of an unusually extensive interface between PDZ_DVL2_ and its cognate atypical PBMs from Daple or Girdin-L.

Next, we used isothermal calorimetry (ITC), titrating 40 μM Lip-PDZ_DVL2_ with1 mM Lip-PBM_Daple_ or 1.33 mM Lip-PBM_FZD4_, which allowed us to determine an apparent *K*_D_ of 9.1 ± 0.1 μM for Daple (**Fig. 3A**). By contrast, binding of Lip-PBM_FZD4_ to Lip-PDZ_DVL2_ is not measurable by ITC under these conditions (**Fig. 3B**). This indicated that the affinity of PDZ_DVL2_ to this canonical class I PBM is at least one order of magnitude lower than its affinity to the atypical PBM-like sequence of Daple. Taken together with our NMR results, this demonstrates that these atypical PBMs in the C-termni of Daple and Girdin-L interact with PDZ_DVL2_ far more extensively than canonical PBMs (*14*, *15*, *27*) and with unusually high affinity.

**Fig. 3.**
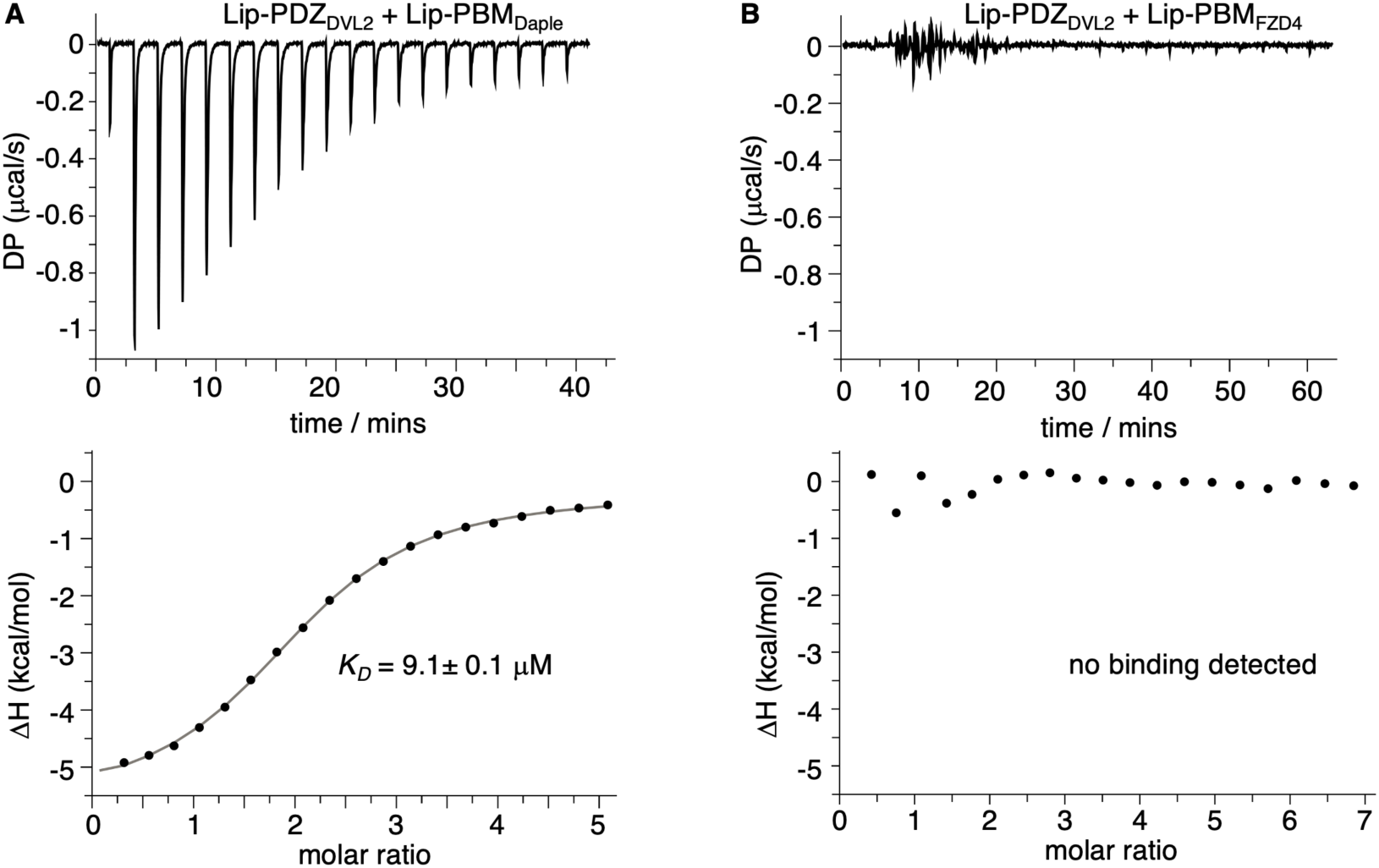
The Daple PBM binds to PDZ_DVL2_ with high affinity. ITC profiles of 40 μM Lip-PDZ_DVL2_ titrated with (**A**) 1 mM Lip-PBM_Daple_ or (**B**) 1.33 mM Lip-PBM_FZD4_; in each panel, the *K*_D_ value was calculated from n=3 independent experiments, and its SD is given. The non-sigmoidal curve and large standard error of fit in the case of Lip-PBM_FZD4_ (**B**) indicates that its binding to Lip-PDZ_DVL2_ is not detectable.

### PDZ_DVL2_ undergoes a structural change upon binding to the Daple PBM

To examine these atypical PBM interactions at the structural level, we attempted to co-crystallize synthetic Daple or Girdin PBM peptides with purified PDZ_DVL2_, but this proved challenging because of the low solubility of these peptides. We therefore resorted to linking them to the C-terminus of PDZ_DVL2_ through a flexible linker as previously published (*26*). Multiple linkers and peptide lengths were tested, but only one combinations produced diffracting crystals. This allowed us to determine the high-resolution structure of the Daple PBM-like sequence (PBM_Daple_) in complex with PDZ_DVL2_ and to compare this to a similarly determined structure of a PDZ_DVL2_ complex with bound PBM from FZD4 (PBM_FZD4_), and also to the structure of unliganded PDZ_DVL2_ (solved at 1.40, 1.75 and 1.49 Å, respectively; **Table S2**).

As expected, both PBM_Daple_ and PBM_FZD4_ were bound to their cognate cleft of PDZ_DVL2_, which is largely formed by α2 and β2 (*21*) (**Fig. 4A**), but they displayed distinct binding modes. Consistent with our NMR results, the contacts of the canonical PBM_FZD4_ with PDZ_DVL2_ were limited to its C-terminal three residues (which conform to a class I PBM, Ser/Thr-X-Φ), with the C-terminal valine (Val^537^, at position 0) buried in the hydrophobic anchoring pocket of the PBM cleft of PDZ_DVL2_ (**Fig. 4B**). Therefore, the interaction mode of PBM_FZD4_ with its cognate cleft in PDZ_DVL2_ followed that of other class I-III PBMs (*14*, *15*).

**Fig. 4.**
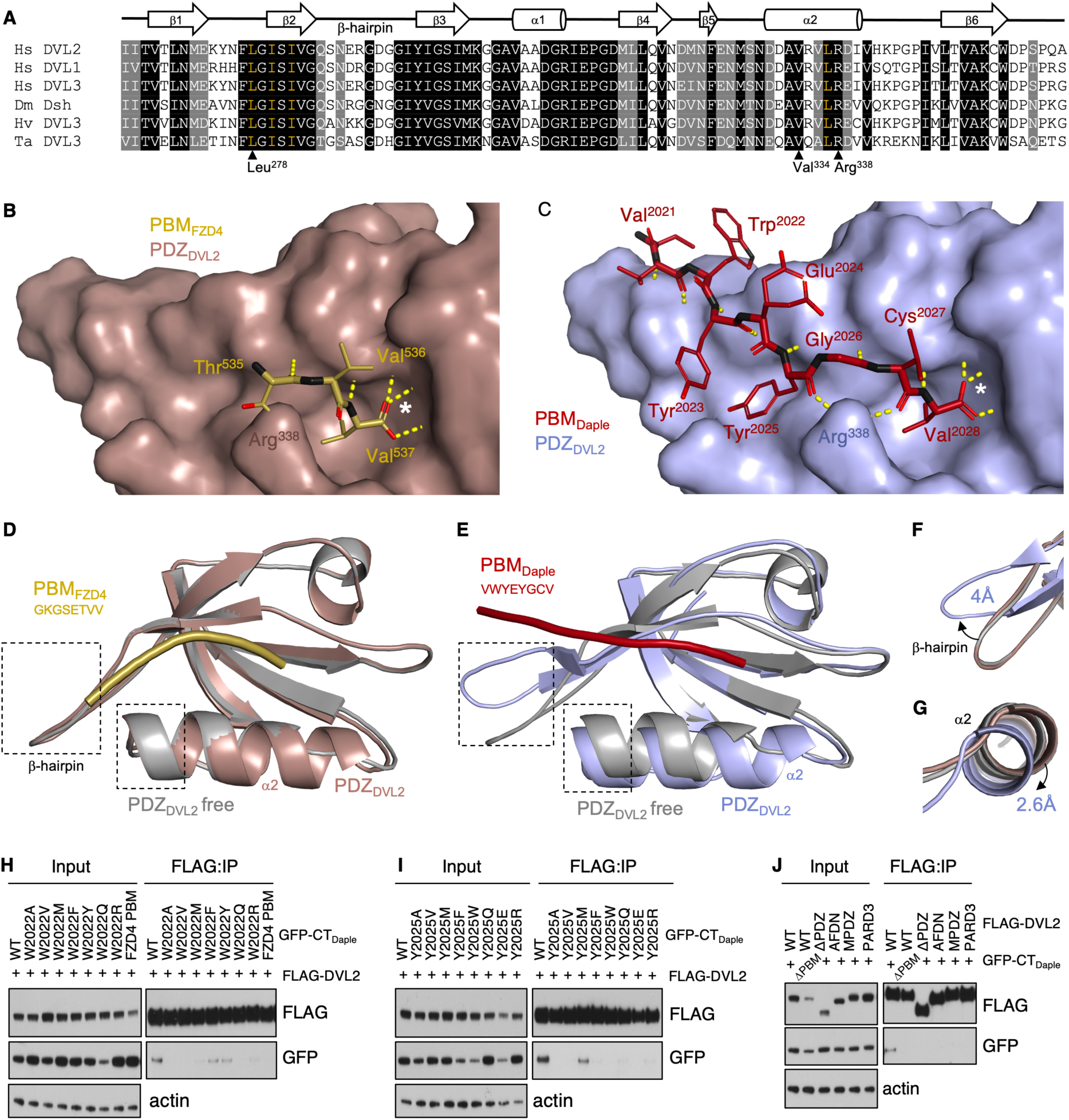
The binding of PBM_Daple_ to PDZ_DVL2_ triggers a structural change. (**A**) Sequence alignment of PDZ domains from diverse species (Hs, *Homo sapiens*; Dm, *Drosophila melanogaster*; Hv, *Hydra vulgaris*; Ta, *Trichoplax adhaerens*); indicated are secondary structure elements (*above*), conserved residues (*highlighted*) and key residues of the PBM-binding cleft (*beneath*); *yellow*, residues of the hydrophobic anchoring pocket. (**B-E**) Crystal structures of (**B, D**) PBM_FZD4_ (*gold*) in complex with PDZ_DVL2_ (*wheat*) or (**C, E**) PBM_Daple_ (*maroon*) in complex with PDZ_DVL2_ (*light blue*); (**B, C**) peptides in stick representations, PDZ_DVL2_ in surface representations; *white asterisks*, hydrophobic anchoring pockets of PBM-binding cleft; *yellow dashed lines*, backbone hydrogen bonds; (**D, E**) PBM-bound structures were superimposed on the structure of unliganded PDZ_DVL2_ (*gray*). (**F, G**) Positional shifts of the (**F**) β-hairpin and (**G**) α2 of PDZ_DVL2_ induced by its binding to PBM_Daple_. (**H-J**) Western blots of co-IPs of WT or mutant GFP-CT_Daple_ with FLAG-DVL2 after co-expression in HEK293T cells, probed with antibodies as indicated on the right.

By contrast, PBM_Daple_ adopted a β-strand conformation, with each of the C-terminal eight residues displaying typical patterns of β-strand hydrogen bonding with residues forming the PDZ_DVL2_ cleft (**Fig. 4C**). The side chain of the C-terminal valine (Val^2028^, at position 0) was buried in the hydrophobic anchoring pocket (formed by Leu^278^, Ile^280^, Ile^282^ and Leu^337^ of PDZ_DVL2_; **Fig. 4A**, *yellow*), and its carboxylate group formed hydrogen bonds with the main chain amides of the carboxylate-binding loop of PDZ_DVL2_ (of Gly^279^ and Ile^280^; **Fig. 4C**). In addition, the backbone carbonyls of the cysteine (Cys^2027^, at position -1) and tyrosine (Tyr^2025^, at position -3) formed hydrogen bonds with the side chains of Arg^338^ of PDZ_DVL2_, conforming to the canonical mode of PDZ-PBM binding (*24*), but distinct to the binding mode between PDZ_DVL2_ and PBM_Daple_ previously proposed based on structural modeling (*37*). However, Gly^2026^ at position -2 of the Daple PBM adopted a positive φ angle and, consequently, the direction of the side chain of the preceding Tyr^2025^ was flipped so that this Tyr-Gly pair occupied position -2 within the PDZ cleft. This shift in register was continued to its position -3 which was occupied by Glu^2024^.

Of note, while the structure of PDZ_DVL2_ is unaffected by its binding to PBM_FZD4_ (**Fig. 4D**), the binding of PBM_Daple_ to PDZ_DVL2_ induced a substantial structural change of this PDZ domain (**Fig. 4E**): its β-hairpin involving Glu^288^ was displaced by 4 Å from its position in the unliganded domain (**Fig. 4F**) and its α2 helix was displaced by 2.6 Å (**Fig. 4G**), although the position of the β2 strand was mostly unchanged. This ligand-induced structural change appears unique amongst PDZ-PBM interactions (*14*, *15*, *24*, *27*) and may reflect the unusually extensive interaction between PBM_Daple_ and PDZ_DVL2_. It could also provide an explanation for the unprecedented high affinity between PBM_Daple_ and PDZ_DVL2_ (**Fig. 3A**) which allowed detection of this interaction by coIP assays (**Fig. 1D-F**).

We noticed that a similarly extensive PDZ_DVL2_ binding mode and shift in register was previously reported for an artificial peptide (called C1) found in a high-throughput phage-display screen of a large random peptide library (*26*). Indeed, this C1 peptide shows sequence similarity to PBM_Daple_ in that both peptides have Trp at position -6 and Tyr-Gly at positions -2 and -3. To determine the functional relevance of the aromatic residues at positions -3 (Tyr^2025^) and -6 (Trp^2022^), we substituted them with different amino acids and tested these substitutions in coIP assays. This revealed that only aromatic substitutions (W2022F and W2022Y) were tolerated at position -6 (**Fig. 4H**) while only methionine (Y2025M) was tolerated at position -3 without causing a substantial reduction of the coIP signals (**Fig. 4I**). As expected, a substitution of the Daple C-terminus with the PBM of FZD4 also failed to coIP with PDZ_DVL2_ (**Fig. 4H**). Taken together, these results provide compelling evidence that the aromatic residue at position -6, upstream of the three residues of a typical PBM, contributes to PDZ binding and to its specificity for PDZ_DVL2_.

Finally, we tested whether other PDZ domains implicated in binding to Daple or Girdin-L (*28*, *42*, *43*) could substitute for PDZ_DVL2_ in this coIP assay. We replaced PDZ_DVL2_ in the context of full-length FLAG-DVL2 with the PDZ domain from Afadin (AFDN), or with the third PDZ domain from the Multiple PDZ domain (MPDZ) subunit of the Crumbs cell polarity complex, or from the cell polarity protein Par-3 (PARD3). However, none of the three DVL2 chimera coIPed robustly with GFP-DapleC_min_, in contrast to WT FLAG-DVL2 (**Fig. 4J**). This further consolidates the notion that the extended ligand-binding cleft of PDZ_DVL2_ is uniquely geared to its strong and specific interaction with Daple or Girdin-L.

### Girdin-L is required for cell motility and disassembly of primary cilia

Previous studies have revealed that Daple and Girdin-L rely on their C-termini for their associations with apical cell junctions (*28*, *43*) and for various cellular functions such as Wnt5a-dependent cell migration and activation of Rac (*16*), facilitating actomyosin-driven apical constrictions of neuroepithelial cells (*28*, *43*), and positioning basal bodies of motile cilia in ciliated cells of the nervous system (*44*, *45*). However, most of these studies were based on assaying overexpressed Daple or Girdin mutants in the presence of endogenous WT protein which may have complemented their functional defects by oligomerizing with these mutants, which could have masked their functions. Functional redundancy between these paralogs may have further confounded the interpretation of the results from these analyses.

We therefore set out to investigate these functions systematically, by using CRISPR engineering of the genes encoding Daple and Girdin in HEK293T cells to generate double null mutants for both (*DG^DKO^*). We also generated a PBM deletion mutants of Girdin-L (by targeting introns 31-32 to delete the PBM-containing C-terminus of Girdin-L while leaving its shorter isoform intact) in a Daple null mutant background (*D^KO^; G^ΔPBM^*). In each case, multiple independent cell lines were isolated for phenotypic analysis of growth, migration and ciliogenesis (see **Fig. 5A**, for a summary of our results).

**Fig. 5.**
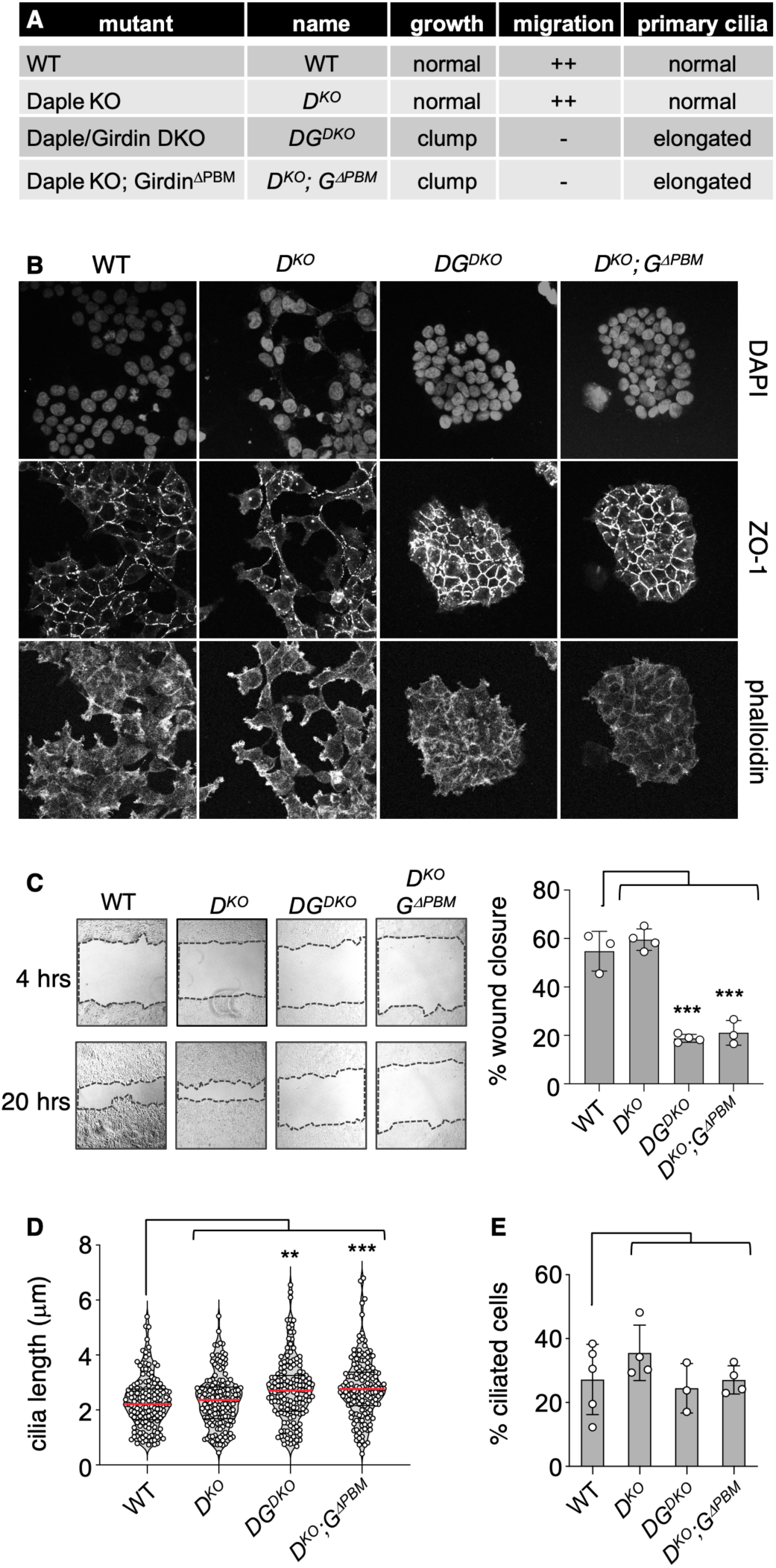
Cell morphology and motility depend on PDZ_DVL2_ and its cognate PBM of Girdin-L. (**A**) Table summarizing CRISPR/Cas9-edited cell lines and their associated phenotypes. (**B**) Confocal images of different HEK293T cell lines (as indicated above panels) fixed and probed with antibodies as indicated on the right. (**C**) Migration of WT and mutant HEK293T cells in scratch assays, as indicated; *left*, representative cell cultures imaged 4 or 20 hrs after scratching; *right*, error bars representing SD of n≥3 independent experiments, each experiment including *n=6* technical repeats; one-way ANOVA with Dunnett’s post hoc; ***=*P*<0.001. (**D**) Violin plots of ciliary lengths in confocal images stained for cilia (ARL13B, *red*) and centrioles (pericentrin, *green*); n≥30 primary cilia were measured in each of >3 independent experiments; red line, median; one-way ANOVA with Dunnett’s post hoc; **=*P*<0.01, ***=*P*<0.001. (**E**) Percentage of ciliated cells; graph plots mean of >3 independent experiments; error bars, SD; one-way ANOVA with Dunnett’s post hoc.

Both types of Girdin mutants (*DG^DKO^*, and *D^KO^; G^ΔPBM^*) showed a pronounced tendency to grow as dense colonies or ‘clumps’. We therefore seeded these mutant cells at a low confluence and allowed them to grow for several days before fixing and staining them for F-actin and zonula occludens-1 (ZO-1), a component of apicolateral tight junctions. In each case, we observed fewer lamellipodial projections of the mutant cells and increased ZO-1 expression at their tight junctions compared to their *D^KO^* or WT parental controls (**Fig. 5B**). The clumping phenotype also correlated with reduced cell migration in ‘scratch’ assays, whereby the rate of migration of the mutant cells was reduced to ∼30-50% of that of their parental controls (**Fig. 5C**).

We also analyzed the primary cilia of these mutants, by co-staining cells for the ciliary marker ARL-13B and the centriolar marker pericentrin and measuring the ciliary length of >30 mutant cells in at least three batches of independently cultured cells. In each case, a substantial proportion of the mutant cells exhibited elongated cilia compared to their parental controls (**Fig. 5D**) although the percentage of ciliated cells was unchanged in the mutant lines (**Fig. 5E**). This suggested that Girdin-L relies on its atypical PBM to promote cell motility and to reduce the length of primary cilia, by antagonizing their assembly or promoting their disassembly. Since these mutant phenotypes were not observed in *D^KO^* cells, this indicates redundancy between Daple and Girdin-L with regard of these functions.

### Ciliary length depends on PDZ_DVL2_ and its cognate PBMs of Daple and Girdin-L

To test whether the ciliary defect in the *D^KO^; G^ΔPBM^* mutants was due to an interaction of Daple and Girdin-L PBMs with PDZ_DVL2_, we generated three lines bearing deletions of the PMBs of Daple and Girdin-L (*DG^ΔPBM^)* and examined their primary cilia alongside DVL-null mutant HEK293T cells (lacking DVL1-3, *DVL^TKO^*) whose Wnt response can be restored by stably re-expressing WT DVL2-GFP or mutant DVL2^ΔPDZ^ at physiological levels (*14*). The primary cilia in *DG^ΔPBM^*mutant cells were significantly longer than those of parental control cells (**Fig. 6A-D**), in some cases up to ∼4x their normal length (**Fig. S4A**). We also noticed persistence of primary cilia in mitotic cells (as determined by condensed chromatin visible by DAPI; **Fig. S4B**) whose disassembly prior to M phase (*46*) was evidently delayed or blocked. *DVL^TKO^* cells also showed elongated cilia (**Fig. 6E**), a mutant phenotype that was rescued by re-expressed WT DVL2-GFP but neither by the ΔPDZ deletion mutant nor by the V334E cleft mutant (**Fig. 6F-H**). Quantification of >30 primary cilia in three independently cultured batches of cells confirmed that their primary cilia were significantly elongated compared to parental control cells or *DVL^TKO^* cells complemented with WT DVL2-GFP (**Fig. 6I**). The percentage of ciliated cells was normal in each case except for the *DVL^TKO^*cells which exhibited fewer ciliated cells (**Fig. 6J**), consistent with a previously reported role of DVL2 in ciliogenesis (*47*). This role appears to be independent of its PDZ domain since the percentage of ciliated cells was not significantly reduced in *DVL^TKO^* complemented with mutant DVL2^ΔPDZ^-GFP (**Fig. 6J**). This indicated two separate functions of DVL2 with regard to primary cilia, namely a PDZ-independent role in promoting their formation and a distinct role in promoting their disassembly prior to mitosis which depends on its PDZ domain and its cognate PBMs of Daple or Girdin-L.

**Fig. 6.**
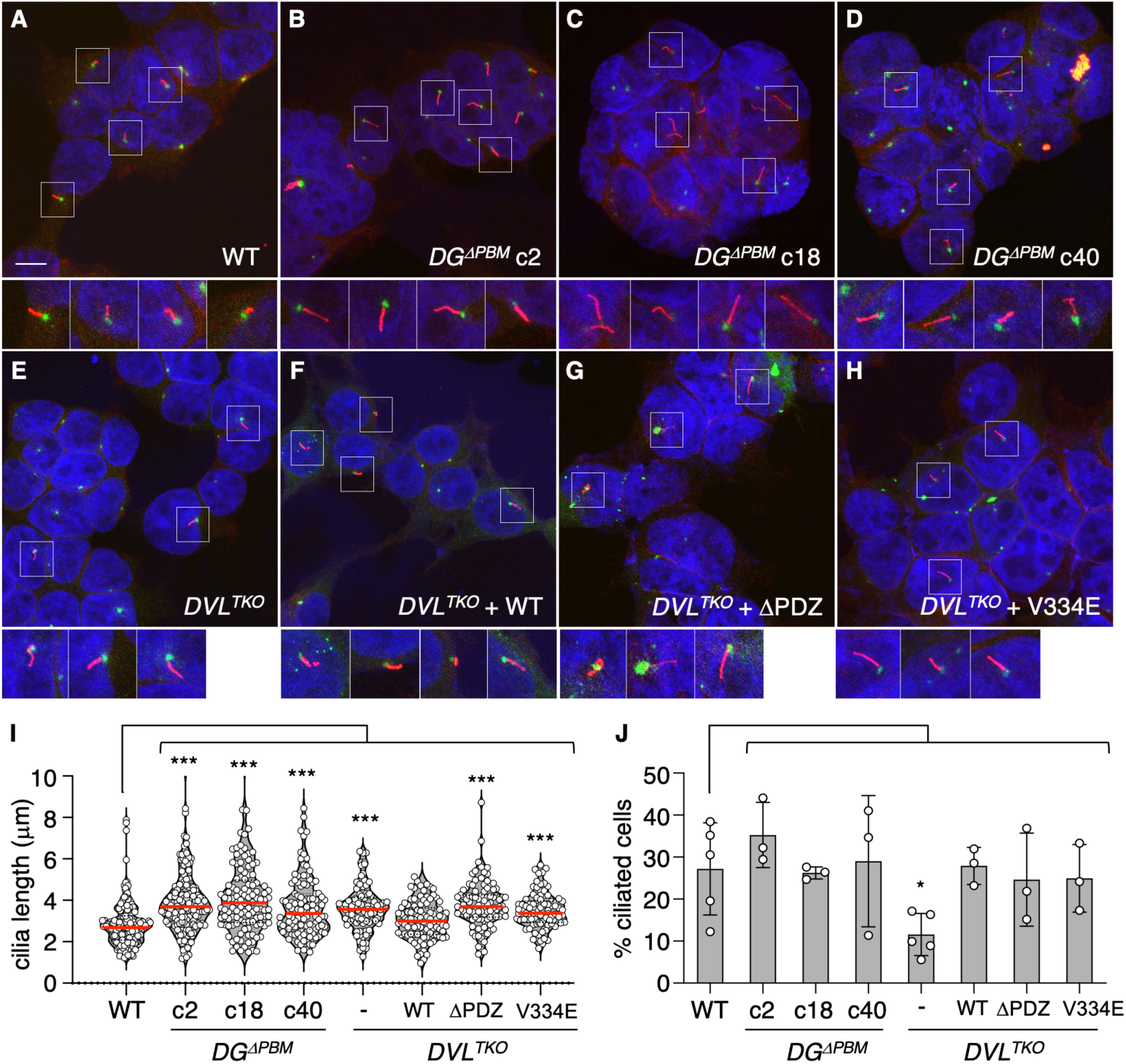
The length of primary cilia depends on PDZ_DVL2_ and its cognate PBMs in Daple and Girdin-L. (**A-H**) Representative merged images of cells, as indicated, stained for ARL13B (*red*), pericentrin (*green*), DAPI (*blue*); scale bar shown in (**A**), 10 μm (representative of all panels); beneath, selected primary cilia at high magnification, box diameter 10 μm; note that the green signals in (**F-H**) represent pericentrin staining superimposed on DVL2-GFP fluorescence. (**I**) Violin plots of ciliary lengths in confocal images; n≥30 primary cilia were measured in each of three independent experiments; red line, median; one-way ANOVA with Dunnett’s post hoc; ***=*P*<0.001. (**J**) Percentage of ciliated cells; graph plots mean of >3 independent experiments; error bars, SD; one-way ANOVA with Dunnett’s post hoc.

### Wnt5a-driven ciliary disassembly depends on PDZ_DVL2_ and its cognate PBMs from Daple or Girdin-L

Primary cilia are not only disassembled prior to mitosis, but also in response to Wnt5a, the non-canonical Wnt that accelerates their disassembly through a Dishevelled-dependent signaling axis (*48*). We therefore tested whether this Wnt5a-dependent process requires Daple and Girdin-L. First, we optimized our conditions with control HEK239T cells, by serum-starving them for 24 hours and subsequently treating with Wnt5a-or mock-conditioned media for 7 hours before fixing and staining them, as previously described (*48*). Under these conditions, we observed a modest reduction of ciliary length upon Wnt5a treatment compared to mock-treated cells (**Fig. 7A**). We next treated our three *DG^ΔPBM^* cell lines with Wnt5a, alongside *DVL^TKO^* cells complemented with various DVL2 mutants. In parental HEK293T cells and in *DVL^TKO^* cells stably re-expressing WT DVL2-GFP but neither ΔPDZ nor V334E mutant, nor in any of the three *DG^ΔPBM^* lines, the length of primary cilia was reduced after stimulation by Wnt5a (**Fig. 7B**), as was the percentage of ciliated cells (**Fig. 7C**). Together with evidence from previous work (*48*), our results support the notion that DVL2 renders primary cilia responsive to Wnt5a by binding to Daple or Girdin-L through its PDZ domain.

**Fig. 7.**
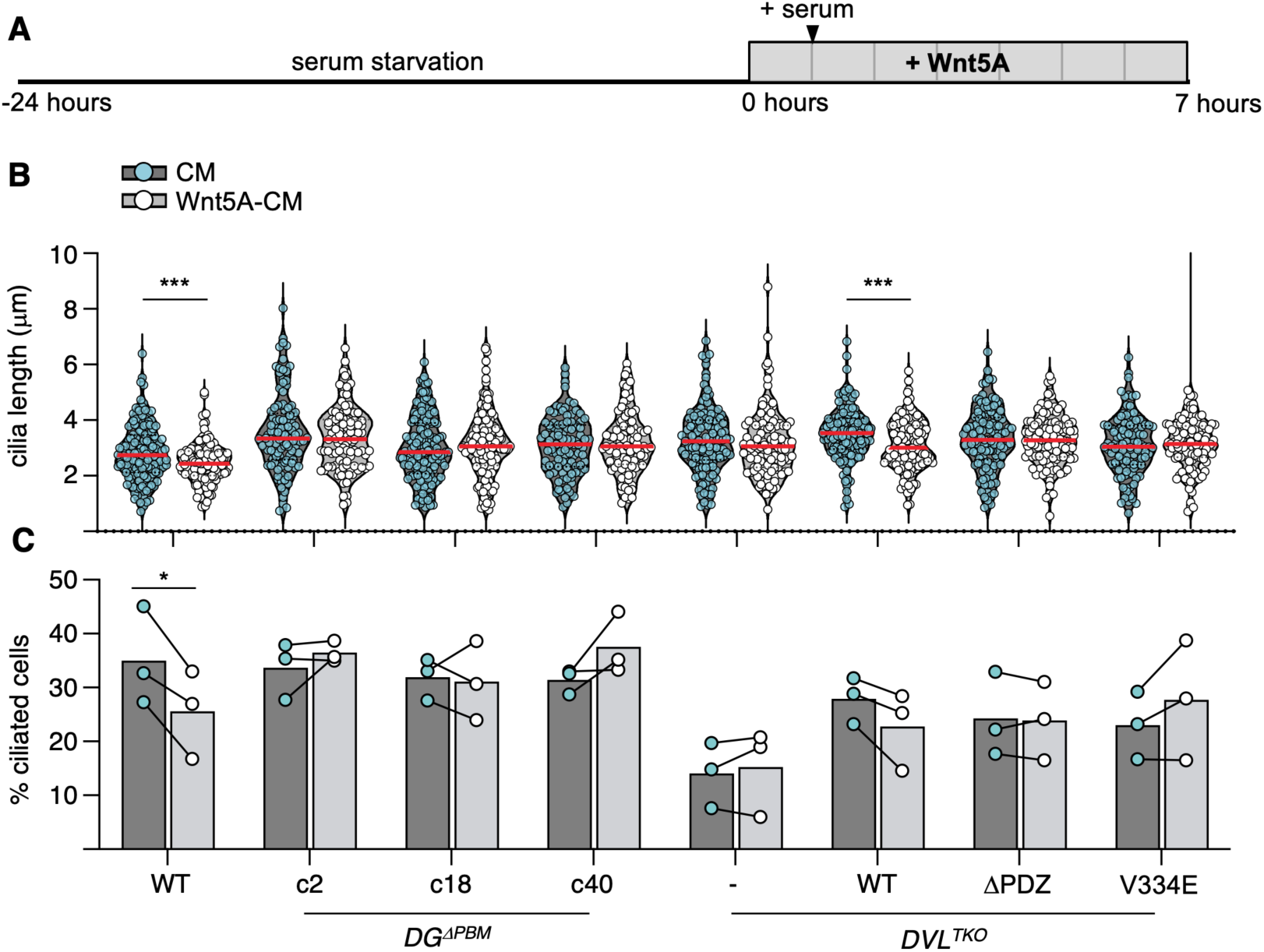
Wnt5a-driven disassembly of primary cilia depends on PDZ_DVL2_ and the PBMs of Daple and Girdin. (**A**) Cartoon of experimental design; HEK293T cells were serum starved for 24 hrs and subsequently treated with Wnt5a-conditioned or control media (CM) for 7 hours (supplemented with serum after one hour), fixed and stained with antibodies against ARL13B and pericentrin. (**B**) Violin plots of ciliary length as in **Figs. 5D & 6I**; n≥30 primary cilia were measured in each of three independent experiments; red line, median; one-way ANOVA with Dunnett’s post hoc; ***=*P*<0.001. (**C**) Bar graph showing percentages of ciliated cells paired from individual experiments with CM-(*blue circle*) or Wnt5a-treated (*white circle*) cells from the same WT and mutant lines shown above (**B**); paired students t-test; *=*P*<0.05.

## DISCUSSION

Our work uncovered a role for the PDZ domain of DVL2 in mediating cell cycle-dependent and Wnt5a-driven disassembly of primary cilia through its binding to Daple or Girdin-L. This was mediated by a deeply conserved eight-residue PBM-like sequence found uniquely at the C-termini of these paralogs that engages in extensive interactions with PDZ_DVL2_. The functional relevance of this interaction was indicated by the requirement of these atypical PBMs and the structural integrity of their cognate binding cleft in PDZ_DVL2_ for the control of ciliary length.

The diversity of PDZ-binding motifs is characterized by their classification into numerous categories, typically consisting of three-residue motifs located at the extreme C-termini of target proteins (*22*). Because of this diversity, a wide array of PBM-bearing proteins are able to bind to PDZ domains (*24*), but not all of these are biologically relevant interactions (*15*). Our study established that the PDZ domain of DVL2 possesses an extended and highly adaptable ligand-binding cleft (*26*) that engages in functionally relevant interactions with all eight residues of the Daple C-terminus, resulting in an unusually high affinity compared to other known PBM-PDZ interactions (*14*, *15*, *26*, *27*).

The same likely applies to the Girdin-L C-terminus, given its extensive interactions with PDZ_DVL2_ (**Fig. 2B, E**) and its C-terminal sequence which is almost identical to that of Daple (**Fig. 1A; Fig. S1**): recall that the only difference between the two is the C-terminal-most residue – valine in Daple, but isoleucine in Girdin-L. Indeed, human Girdin-L seems exceptional in this respect, as Girdin orthologs in orther species exhibit a valine at their C-termini (**Fig. S1**). This C-terminal-most residue is invariably hydrophobic in all classes of PBM as it binds to a hydrophobic pocket within the binding cleft of PDZ domains that may provide anchoring (**Fig. 4B, C**) (*22*, *24*). Similarly to valine, isoleucine has a branched hydrocarbon side chain, but its branch is structurally distinct and contains four rather than three carbons, both of which may affect the strength of its anchoring within, and its affinity to, PDZ_DVL2._ Nevertheless, given the close similarity between the heat-maps produced by PBM_Girdin_ and PBM_Daple_ (**Fig. 2D, E**), it seems likely that both PBMs engage in comparable high-affinity interactions with PDZ_DVL2_ which distinguish them from other ligands of this domain. Of note, not all known functional interactions of PDZ_DVL2_ are mediated by PBM-like sequences, for example a well-known universal feedback regulator of Wnt signaling called Naked Cuticle binds to this domain through its single EGF-hand (*49*, *50*).

How do Daple and Girdin-L function as effectors of DVL2 in the disassembly of primary cilia? In cycling cells, this process occurs shortly before mitosis and involves two steps: first, Aurora A kinase (AurA) is activated through its association with the HEF-1/Nedd9 scaffold, to initiate the depolymerization of microtubules within the axoneme of the cilium, which is followed by the resorption of the ciliary membrane (*46*). The first step can be stimulated by Wnt5a which signals through DVL2, casein kinase 1ε and polo-like kinase1 to stabilize HEF-1 and associated AurA (*48*). Given that Daple and Girdin can serve as adaptors for dynein (*51*), it is possible that these proteins, following their DVL2-dependent recruitment to basal bodies of primary cilia, relocate dynein to the cell cortex where this motor protein cooperates with kinesins (as mentioned earlier) to promote the retrograde transport of ciliary components out of the primary cilium, thereby promoting the depolymerization of ciliary microtubules.

Alternatively, or in addition, it is possible that the effector function of Daple and Girdin involves their activity in promoting G protein exchange through their GBA (**Fig. 1C**), e.g. of Rac1 to promote rearrangement of the cortical actin cytoskeleton and cell motility (*16*, *32*) (**Fig. 5B, C**). Future work will be required to distinguish between these possibilities and to pin down the molecular step(s) during ciliary disassembly governed by the interaction of Daple and Girdin-L with ciliary DVL2 during the normal cell cycle, and how this process is accelerated by Wnt5a.

Defects in primary and motile cilia contribute to a spectrum of disorders collectively known as ciliopathies. Notably, ciliary dysfunction can lead to cerebral anomalies, as demonstrated by the development of hydrocephalus in following conditional depletion of Dvl1-3 or complete knock-out of Daple in mice, owing to defects in motile cilia of ependymal cells (*45*, *52*). In humans, mutations in *CCDC88C* (the gene encoding Daple) have been identified in multiple families with primary non-syndromic congenital hydrocephalus (*53*, *54*). In one specific case, a homozygous mutation caused a frameshift in the last exon of *CCDC88C*, resulting in a premature stop codon that truncated the extreme C-terminus of Daple (*53*), underscoring the pivotal role of its PBM. Furthermore, defects in ciliary disassembly have been implicated in microcephaly and malformations of cortical development (MCD). For instance, disease-causing mutations in key regulators of ciliary disassembly, such as the dynein light-chain protein Tctex-type 1 (or DNYLT1) and kinesin KIF2A have been identified in MCD patients (*55–57*). Furthermore, mutations in *CCDC88A* (the gene encoding Girdin) have been found in patients with progressive encephalopathy characterized by oedema, hypsarrhythmia, and optic atrophy (PEHO)-like syndrome (*58*). Similarly, knock-out of Girdin in mice caused PEHO-like phenotypes, including microcephaly, which indicated a crucial role of Girdin in the brain (*58*). Our discovery that the interaction between PDZ_DVL2_ and its cognate Daple and Girdin-L ligands promote disassembly of primary cilia provides a possible molecular explanation for some of these developmental cerebral anomalies in mice and humans.

## MATERIALS AND METHODS

### Cell cultures and lines

HEK293T (ATCC, Cat#CRL-3216), HEK293T DVL TKO, FlpIn T-REx (Thermo Fisher Scientific, Cat#R78007) and HeLa (ATCC, Cat#CCL-2) cells were cultured in 6-well or 24-well culture dishes in DMEM+GlutaMax (Gibco, Cat#11594446), supplemented with 10% fetal bovine serum (FBS; Gibco, Cat#11594446) plus 1% penicillin/streptomycin (Sigma) at 37°C in a humidified atmosphere with 5% CO_2_, and regularly screened for mycoplasma.

Wnt3a-, Wnt5a-and control conditioned media was collected from L-cells (ATCC, CRL-2648; CRL-2814; CRL-2647) following the manufacturer’s instructions, and added to cells for the time periods indicated in the text and figures. To generate DVL2-expressing HEK293T cell lines for DVL complementation tests, DVL2-GFP was inserted into the pBABE vector and stably re-expressed in DVL TKO, as previously described (*14*).

### Generation of plasmids

Sequences for in vitro and cell-based assays were amplified by polymerase chain reaction from either plasmid templates or synthetic genes (gBlocks, IDT), cloned into mammalian or bacterial expression vectors by restriction-free cloning using Gibson Assembly Master Mix (NEB, Cat#E2611L). Point mutations and deletions were generated by Quikchange, using KOD Hot Start DNA polymerase (Merck Millipore, Cat#71086) or Q5 polymerase (NEB, Cat#M0491L). All plasmids were verified by DNA sequencing.

### CRISPR engineering

The CRISPR design tool CRISPOR (crispor.tefor.net) was used to design the following gRNAs targeting Daple (ACTTTTGGCCCGTTTGGAAG, TGGTCAAATTCTGAATGCGA, GCGGGGATCCGCAGACCGTG) or Girdin (GACCAACCTTGATGAATATG, CTGAGGATAATCAAACTGTT). These were inserted into PX458 (Addgene) by Bbs1 (NEB, Cat#R3539S) cloning. Single cell clones were grown in culture media supplemented with Plasmocin (InvivoGen, Cat#ant-mpp) to protect against mycoplasma infection and analyzed by DNA sequencing. For *DG^DKO^* and *D^KO^; G^ΔPBM^*CRISPR edits were introduced sequentially, first targeting Daple and then Girdin. Briefly, cells were lysed in Squash buffer (10 mM Tris pH8.0, 1 mM EDTA, 25 mM NaCl, supplemented with 200 μg/ml Proteinase K) and heated in a Thermal Cycler (65°C for 15 minutes, 96°C for 2 minutes, 65°C for 4 minutes, 96°C for 1 minute, 65°C for 1 minute, 96°C for 30 seconds). DNA was amplified using the following primers (Daple exon2_ampF: AGATACCAGCGTCTCCACCT; Daple exon2_ampR: GGGAGAAAAGTGGCGAGGAA; Daple exon2_seqR: CTCCTGTCTCGTGTCTGTTGC; Daple exon3_ampF: CACAGCCTCATGGTCACCTA; Daple exon3_ampR: CACGATGCTACACACAGCTC; Daple exon3_seqF: TCAACGCACCATCTCATCCTC; Daple exon30_ampF: CTCCTGCACTTCTCACCTGC; Daple exon30_ampR: GTCAGAAAGCCCACTCCCTC; Daple exon30_seqR: CGTGTTTTCTGTGTCTGCCG; Girdin exon2_ampF: GGCCAATGAACCAAATTAGGGGC; Girdin exon2_ampR: CCCCTCCTCCACGATGGGAT; Girdin exon2_seqR: GGACTTTCATTGATTTCTAC; Girdin exon31_ampF: TGGGTTACCTCCTAGGCCAG; Girdin exon31_ampR: ACCACACCAAAACCTGCATG; Girdin exon31_seqR: TGTTGCACAGTGTTCAGCTG) and sequence chromatograms were imported into TIDE (tide.nki.ni).

### coIP assays

HEK293T cells were seeded at ∼70% confluency and transfected with a 1:3 mixture of DNA:PEI (polyethylenimine, Polysciences, Cat#23966, according to manufacturer’s instructions) after cells had attached. For all coIPs, one well of a 6-well plate per coIP was used. Cells were lysed ∼24 hours post-transfection in 20 mM Tris pH 7.4, 200 mM NaCl, 10% glycerol, PhosSTOP (Sigma, Cat#04906837001), EDTA-free protease inhibitor cocktail (Roche, Cat#04693159001) and 0.1% Triton-X. Lysates were sonicated and cleared by centrifugation (16,100 *g*, 10 minutes), and supernatants were incubated with GFP-trap (Chromotek, RRID: AB_2631357) or α-FLAG M2 agarose beads (Sigma, RRID: AB_2637089) for >90 minutes at 4°C on an over-head tumbler. Immunoprecipitates were washed three times in lysis buffer and eluted by boiling in 4x lithium dodecyl sulphate (LDS) sample buffer (Invitrogen, Cat#NP0007) for 10 minutes. Input lysates (1% of total) and coIP eluates (20% of total) were separated by SDS polyacrylamide gel electrophoresis (SDS-PAGE), blotted onto polyvinylidene difluoride membranes, checked for equal loading by Ponceau staining and processed for Western blotting using the following primary antibodies: α-GFP and α-FLAG (RRIDs: AB_439690; AB_439687, Sigma), α-actin and α-Girdin (RRIDs: AB_2305186; AB_10859854, Abcam), α-GSK3β (RRID: AB_490890, Cell Signaling Technologies). Primary and secondary antibodies were diluted 1:1000-5000 in phosphate-buffered saline (PBS), 0.01% Triton-X and 5% milk powder. Blots were washed with PBS containing 0.01% Triton-X and developed with ECL Western Blotting Detection Reagent.

### Immunofluorescence and measurement of ciliary length

HEK293T or HeLa cells were seeded on coverslips at ∼70% confluency and transfected after cells had attached. Serum starvation was performed by replenishing media with OptiMEM (Thermo Fisher, Cat#31985070) for 24 hours. Cells were fixed in 4% paraformaldehyde (PFA), permeabilized and blocked in 5% Bovine Serum Albumin (BSA), 0.01% Triton-X. Primary and secondary antibodies were diluted 1:50-4000 in blocking solution. FZD recruitment assays were done as previously described (*7*, *8*, *39*). The following primary antibodies were used: α-GFP and α-FLAG (RRIDs: AB_439690; AB_262044, Sigma), α-pericentrin and α-ZO-1 (RRIDs: AB_304461; AB_2909434, Abcam), α-ARL13B (RRID: AB_2060867, Proteintech) and α-SNAP (RRID: AB_10631145). Cells were also stained with phalloidin-546 (Thermo Fisher, Cat#A22283). Confocal images were taken with a Zeiss confocal microscope. To measure the length of primary cilia, Z-stacks were performed and images flattened. Measurements were performed in ImageJ on clearly identifiable primary cilia (typically >1 μm) using ARL13B and pericentrin as markers. >30 primary cilia were measured for each condition, and experiments were repeated >3 times.

### Scratch assays

HEK293T cells were seeded and grown to confluence in 24-well culture dishes and a manual scratch was made with a p1000 pipette tip. Cells were gently washed with 1xPBS and media replenished. Cells were returned to the incubator and imaged at 4 and 20 hours. Areas void of cells were measured in ImageJ. For each condition, 6 technical replicates were averaged, and each experiment was repeated >3 times.

### Mass spectrometry

To generate BioID plasmids, coding sequences of DVL2 and its ΔPDZ mutant version were amplified from a DVL2-GFP encoding pEGFPN-1 plasmid (*14*) by PCR and inserted directly upstream of BirA* in pcDNA5/FRT/TO using Gibson assembly (*59*). To generate stable cell lines, FlpIn T-REx cells were co-trasnfected with DVL2-BirA* pcDNA5/FRT/TO plasmids (WT and ΔPDZ) and pOG44 (Flp recombinase vector) and selected with 250 μg/ml hygromycin B (ThermoFisher, Cat#10687010). For each stably transfected cell line, DVL2 expression was induced with tetracycline (Sigma, Cat#T8032) for 24 hours, and biotin (Sigma, Cat#04906837001) labeling was performed for 12 hours. A total of 1.4-2.1 x 10^8^ adherent cells were grown to full confluence, washed once with phosphate-buffered saline, flash-frozen in liquid nitrogen, and stored at -80 °C for 1-20 days. BioID pull-downs were done using Streptavidin Dynabeads (Invitrogen, Cat#65001) as described (*29*), and proteins were eluted from the beads by boiling for 15 minutes in LDS sample buffer (Invitrogen, Cat#NP0007). All samples were resolved on 4-12% Bis-Tris polyacrylamide gels, and gels were stained with Imperial Protein Stain (ThermoScientific, San Jose CA, USA). Gel slices (2-3 mm) were prepared for mass spectrometric analysis by manual *in situ* digestion with trypsin, and digests were analyzed by nano-scale capillary LC-MS/MS using an Ultimate U3000 HPLC (ThermoScientific Dionex, San Jose, USA). The analytical column outlet was directly interfaced via a nano-flow electrospray ionization source, with a hybrid dual pressure linear ion trap mass spectrometer (Orbitrap Velos, ThermoScientific, San Jose, USA). LC-MS/MS data were searched against a protein database (UniProt KB) using the Mascot search engine program (Matrix Science, London, UK). MS/MS data were validated using the Scaffold program (Proteome Software Inc., Portland OR, USA) and processed with R package. The raw mass spectrometry proteomics data have been deposited to the ProteomeXchange Consortium (*60*) via the PRIDE partner repository (*61*).

### Protein expression and purification

Recombinant His6x-Lip-PDZ, His6x-Lip-peptides, and His6x-Lip-PDZ-GS linker-peptide fusions were purified from BL21-CodonPlus(DE3)-RIL cells (Agilent) *E. coli* bacterial strains. Bacteria were grown at 37°C in LB media supplemented with appropriate antibiotic to OD_600_ 0.6, then moved to 24°C, followed by induction with 0.4 mM IPTG at OD_600_ 0.8. Bacteria were harvested by centrifugation (8,000 g, 30 minutes); cell pellets were shock-frozen in liquid nitrogen and stored at -80°C until use. Harvested cells expressing the above proteins were resuspended in lysis buffer (25 mM Tris pH 8.0, 200 mM NaCl, 20 mM imidazole pH 8 and EDTA-free protease inhibitor cocktail; Roche) and lysed by sonication (Branson). Lysates were cleared by ultracentrifugation (140,000 *g*, 30 minutes at 4°C) and incubated with Ni-NTA agarose (Qiagen, Cat#30210) and washed with lysis buffer. After extensive washing, samples were eluted with lysis buffer supplemented with 500 mM imidazole and loaded onto a HiLoad 26/600 Superdex 75 pg column (GE Healthcare) equilibrated in 25 mM sodium phosphate pH 6.7 and 150 mM NaCl. Proteins used for crystallization were treated with in-house produced TEV protease (overnight, 4°C, 1:20 molar ratio). The uncleaved proteins and TEV protease were removed during subsequent incubation with Ni-NTA agarose. Each step of the purification was done at 4°C, and protein purity was assessed by SDS-PAGE. All proteins were concentrated, flash frozen in liquid nitrogen and stored at -80°C until use.

### NMR

For NMR spectroscopy, PDZ_DVL2_ was expressed in M9 minimal medium supplemented with antibiotics, trace elements, 25 ml overnight culture and 2 g of ^15^N-H_4_Cl per litre of expression culture. Cultures were grown and processed essentially as described above. All NMR spectra were recorded using Bruker Avance III spectrometers operating at 600 MHz ^1^H frequency, with 5 mm inverse-detect cryogenic probes and a sample temperature of 298 K using unmodified Bruker pulse programs. Backbone resonance assignments were previously obtained (*14*, *15*). Frequencies were referenced according to the unified scale, with the ^1^H signal of internal dimethylsilapentane sulfonate (DSS) at 0.0 ppm. All spectra were processed with TopSpin version 3 (Bruker) and analyzed using NMRFAM-Sparky version 1.3 (*62*).

Sequence-specific connectivity was aided with the program MARS version 1.2 (*63*). Protein-protein interactions were inferred from chemical shift perturbation or peak height attenuation (‘bleaching’) in 2D fHSQC H_N_-N correlation of 100 µM ^15^N-labeled PDZ_DVL2_. fHSQC spectra were acquired with 256 and 1024 points in *t*_1_ and *t*_2_, respectively, 16 scans per *t*_1_ increment and a recycle delay of 0.9 seconds. Peak heights (*I*) were fit in Sparky. Relative peak height attenuation was calculated from 100x(*I*_ref_–*I*_complex_)/*I*_ref_.

### ITC

Affinities between Lip-PDZ_DVL2_ and Lip-PBM_FZD4_ or Lip-PBM_Daple_ peptide were determined by ITC at 25°C with a Malvern P analytical ITC200 instrument in 25 mM sodium phosphate pH 6.7 and 150 mM NaCl buffers. The concentration of Lip-PDZ_DVL2_ in the cell was 40 μM, and the peptide concentration in the syringe was 1.00 mM and 1.33 mM for Lip-PBM_Daple_ and Lip-PBM_FZD4_ peptide, respectively. Titrations consisted of 19 injections of 2 μL preceded by a small 0.5 μL pre-injection that was not used during curve fitting. Experiments were performed at a reference power of 6 μcal/second and with injections at 180 second intervals with constant stirring at 750 rpm. All ITC binding data were corrected with the appropriate control heats of dilution and fitted using the ‘one set of binding sites’ model in MicroCal PEAQ-ITC analysis software (v1.41). Experiments were performed at least 3 times.

### Protein crystallization and data collection

Purified PDZ and PDZ fusion proteins were concentrated with a 3 kD MWCO Vivaspin 20 concentrator (Sartorius) to 20 mg/ml. Crystallization trials with multiple commercial crystallization kits were performed in 96-well sitting-drop vapor diffusion plates (Molecular Dimensions) at 18°C and set up with a mosquito HTS robot (TTP Labtech). Drop ratios of 0.2 μL + 0.2 μL (protein solution + reservoir solution) were used for coarse and fine screening. Initial hits were obtained under multiple conditions and optimized subsequently. Data for unliganded PDZ were collected from crystals harvested from 0.1 M sodium acetate, pH 4.5, 1 M ammonium sulfate. Data for PDZ-Daple fusion were collected from crystals harvested from 0.1 M sodium acetate, pH 4.6, 3% 2-methyl-2,4-pentanediol. Data for PDZ-FZD4 fusion were collected from crystals harvested from 0.1 M magnesium formate, 15% w/v PEG 3350. Crystals were directly flash frozen in liquid nitrogen after a brief soak in mother liquier supplemented with 25% glycerol. Diffraction data were collected at the Diamond Light Source (DLS, UK) on beamline I04 and I24 . Data processing was performed with XIA2 DIALS and scaled using Aimless from CCP4 (Collaborative Computational Project, Number 4, 1994) (*64*). Structures were solved by molecular replacement using a previously published PDZ structure (PDB code 3CBX). Structure refinement was performed with REFMAC followed by manual examination and rebuilding of the refined coordinates in the program COOT (*65*). Color figures were prepared with PyMOL (Schrödinger).

### Statistical analysis

All error bars are represented as mean ± SD for 3-4 independent experiments. Statistical significance was calculated in Prism V10.0 (GraphPad) by one-way ANOVA followed by Dunnett’s multiple comparisons; *P*-values between indicated data points are given in each figure legend (and are denoted in individual panels as *, *P* < 0.05; **, *P* < 0.01; ***, *P* < 0.001; ****, *P* < 0.0001).

## Acknowledgements

We thank Akira Kikuchi (Osaka University) for the Daple plasmid, Jerome Boulanger (LMB) and the LMB mass spectrometry and cell sorting facilities for expert technical assistance.

## Author Contributions

M. V. G. and M. B. designed the project; M. R, G. B., T. R and M. V. G. performed experiments and analyzed data; M. V. G. and M. B. supervised the study and wrote the manuscript with input from all authors.

## Data and materials availability

Atomic structures and structure factors will be deposited in the Protein Data Bank (PDB). NMR assignment has been deposited in the Biological Magnetic Resonance Bank (https://bmrb.io) under accession number 53371. The mass spectrometry proteomics data will be deposited to the ProteomeXchange Consortium via the PRIDE partner repository. The plasmids are freely available from M. V. G.

