## Supplemental figures and tables for "Dishevelled drives Wnt-stimulated disassembly of primary cilia through a unique PDZ-mediated binding mode with Daple and Girdin"

```
consurf
```

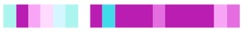

```
Hs Daple           ---LPAKSPGRSPDLAPHLGRAL---EDCSRGSVSKSS-PASPEPGGDPQTVWYEYGCV 2028
Hs Girdin.         ----------------------------------------------------------- 1870
Hs Girdin X1*      SNLQFSINSNN-LVTSACL-------YPDDTEAAL--------LVSEDNQTVWYEYGCI 2027
Mm Daple           ---LLTKSPGRSSDLPPHVKRGP---DDFSQGSSSKST-PASPEPGGDPQTVWYEYGCV 2009
Xl Daple           ---LASAGPGKSRSTSPDCSPAPVCEEEPNKSETLKST-PASPDPSADPQTVWYEYGCV 2058
Xl Girdin          --------PPVTKSDIPNF-PKDDCECPVVQGPSDKET-EISLSVTEDNQSIWYEYGCV 2005
Dr Daple           -----------REAESPLLKKADTTNLSYASKEQPTSK-PASPDPNNDPQTVWYEYGCV 2000
Dr Girdin          ----------------------------------------------------------- 1603
Dr Girdin X1*      --------------------------RPADPEA-----------CVSAEDGLWYEYGCV 1843
Ap Girdin-like     TRDRSVSEPAKAKTGTPT---------KTEQGPGQEEGQNGENKGAKKKSSVWYEYGCV 1962
Ap Girdin          ------------------------------------------DKDSKTGNQVWYEYGCV 1182
Am Daple           ------------------------------------------DKDSKTGNQVWYEYGCV 1258
Dp Hook            ------------------------------EASRDKSVNRKDVNESSKETAIWYEYGCV 1360
My Girdin          -------MGGMSRTVNPGVKSSPSS---TSAAPRPM--KAEENATEPNKNALWYEYGCV 2030
Dm Girdin          ------------------------------------S-A--RKRTEDKSNSIWYEYGCV 1381
Hv Girdin-like     --------INRNKHKSLNL----------ENSDEHQSSFRRSDSTRQSKRKTWYEYGNV 2131
```

**Supplementary Fig. 1. Evolutionary conservation of Daple and Girdin C-termini.** Sequence alignment of Daple and Girdin orthologs (Hs, *Homo sapiens*; Mm, *Mus musculus*; Xl, *Xenopus laevis*; Dr, *Denio rerio*; Ap, *Acanthaster planci*; Am, *Apis mellifera*; Dp, *Danaus plexippus*; My, *Mizuhopecten yessoensis*; Dm, *Drosophila melanogaster*; Hv, *Hydra vulgaris*) indicating conservation of residues in their C-termini, which bear PBM-like motifs; *above*, Consurf plot of conservation across mammals with pink highly conserved and blue not conserved; * indicates splice isoforms of Girdin, which possess PBM-like motifs (in the human isoform in intron 31-32).

**Renko et al., Supplementary Figure 1**

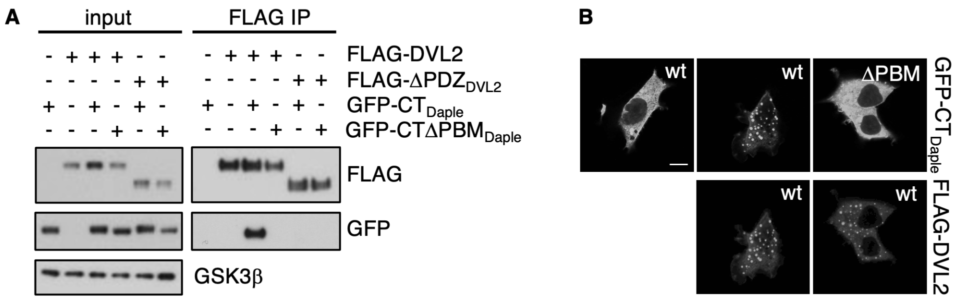

**Supplementary Fig. 2. The PBM-like motif of Daple binds to PDZ$_{DVL2}$.** (**A**) Western blots of coIPs of WT or ΔPDZ mutant FLAG-tagged DVL2 after co-expression with WT or PBM-deleted GFP-tagged mouse Daple C-terminus (GFP-CT$_{Daple}$) in transiently transfected HEK293T cells (as in **Fig. 1F**), probed with antibodies as indicated on the right; representative of n=3 biological repeats. (**B**) Confocal images of representative HEK293T cells (fixed 16 hours after transfection), co-expressing WT or GFP-CTΔPBM$_{Daple}$ and FLAG-DVL2 as indicated; scale bar 10 µm (in all panels); images representative of *n>100* cells visually scored.

**Renko et al., Supplementary Figure 2**

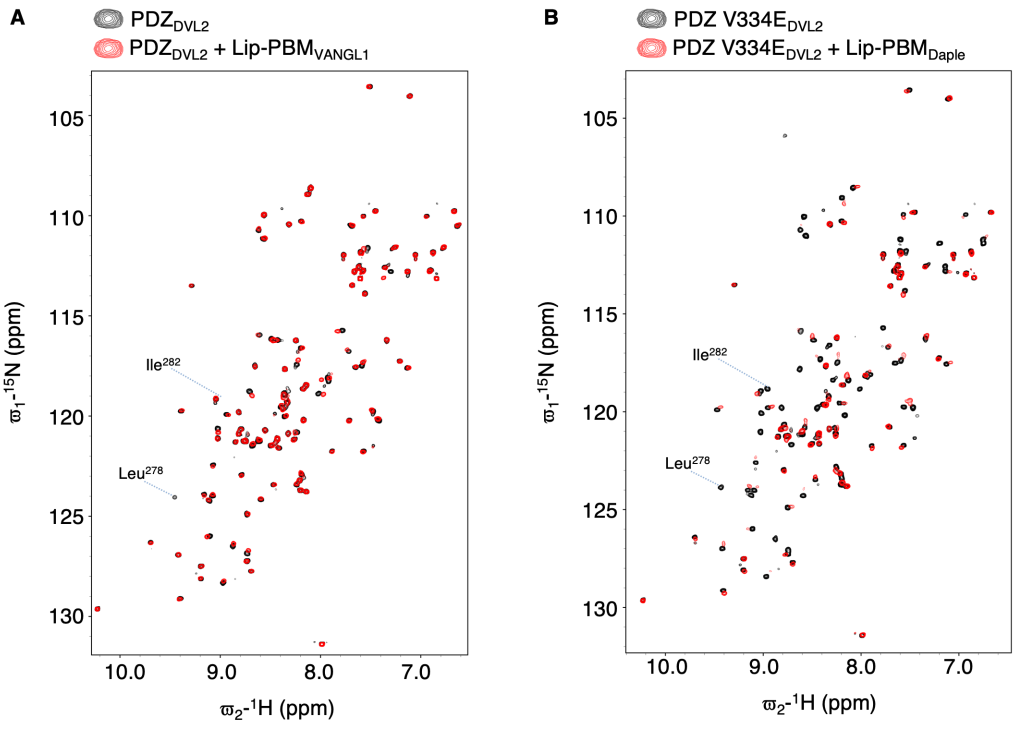

**Supplementary Fig. 3. The binding of PBM$_{Daple}$ to PDZ$_{DVL2}$ is not blocked by the V334E cleft mutation.** Overlays of HSQC spectra of 100 μM WT (**A**) or V334E (**B**) $^{15}$N-PDZ$_{DVL2}$ alone (*black*), incubated with (**A**) 300 μM Lip-PBM$_{VANGL1}$ Or (**B**) 150 μM Daple (*red*).

**Renko et al., Supplementary Figure 3**

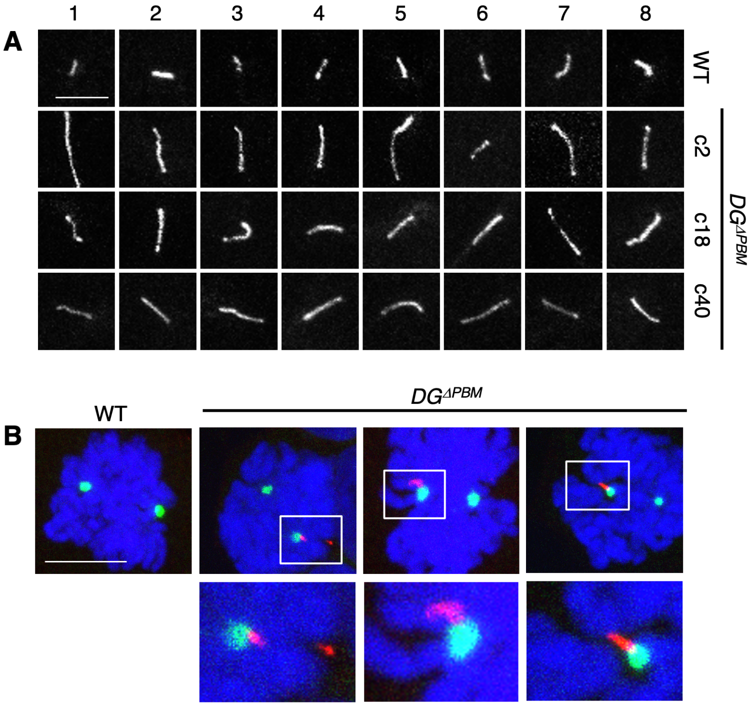

**Supplementary Fig. 4. The length of primary cilia depends on $PDZ_{DVL2}$ and its cognate PBMs in Daple and Girdin-L.** Representative examples of cilia (**A**) at high magnification (*grayscale)* or (**B**) in mitotic cells stained with ARL13B (*red*), pericentrin (*green*) and DAPI (*blue*). Scale bars, 10μm; the mitotic cells in (**B**) were identified on the basis of DAPI staining.

**Renko et al., Supplementary Figure 4**

| | WT/ΔPDZ | | | |
|---|---|---|---|---|
| **Protein** | **expt1** | **expt2** | **expt3** | ***P*-value** |
| CCDC88A | 51/1 | 26/1 | 19/1 | *** |
| CCDC88C | 104/4 | 63/8 | 67/2 | ** |
| CEP192 | 58/3 | 38/7 | 20/3 | ** |
| PSMD8 | 11/2 | 10/2 | 4/1 | *** |
| ALMS1 | 38/5 | 27/10 | 29/6 | ** |
| LDH | 7/1 | 6/1 | 7/4 | * |
| KANK2 | 22/5 | 17/7 | 6/1 | ** |
| RPL7 | 5/1 | 6/1 | 7/4 | * |
| FOXK1 | 4/1 | 2/1 | 10/2 | * |
| PRPF8 | 50/10 | 23/10 | 31/9 | ** |
| hCG_1993905 | 5/1 | 3/1 | 2/1 | * |
| SF3A3 | 3/1 | 5/1 | 2/1 | * |
| SQSTM1 | 10/2 | 6/2 | 4/2 | * |
| ABCF1 | 4/2 | 3/1 | 4/1 | ** |
| CHMP4B | 3/1 | 4/1 | 2/1 | ** |
| VANGL1 | 15/5 | 15/8 | 4/1 | * |
| RPL7A | 9/4 | 12/3 | 8/4 | * |
| SREK1IP1 | 3/1 | 3/1 | 2/1 | ** |
| RPL18A | 6/2 | 7/4 | 3/1 | ** |
| RPL8 | 7/3 | 8/2 | 8/5 | * |
| NSA2 | 7/4 | 6/2 | 5/2 | ** |
| TCP1 | 6/2 | 10/4 | 12/9 | * |
| CCT3 | 6/4 | 6/2 | 13/6 | * |
| OCRL | 23/8 | 14/7 | 5/3 | * |
| SCYL2 | 19/7 | 8/4 | 3/2 | * |
| FLII | 2/3 | 1/2 | 1/4 | * |
| LIMD1 | 1/4 | 3/6 | 1/2 | * |
| EPRS | 1/5 | 3/10 | 8/11 | * |
| DDX20 | 1/5 | 1/4 | 1/3 | *** |
| EPS15 | 1/3 | 1/7 | 1/4 | ** |
| DVL2 (Bait) | 35/35 | 36/35 | 28/24 | ns |

**Supplementary Table 1. List of significant $PDZ_{DVL2}$-sensitive BioID hits**
Numbers represent WT/ΔPDZ ratios of unweighted exclusive unique spectral counts (>95% probability); CCDC88A, Girdin; CCDC88C, Daple. P-values generated by students paired t-test relative to bait, *=P<0.05, **=*P*<0.01, ***=*P*<0.001.

| | $PDZ_{DVL2}$ Free | $PDZ_{DVL2}$ FZD4 | $PDZ_{DVL2}$ Daple |
|---|---|---|---|
| **PDB ID** | | | 9SSJ |
| **Resolution range** | 45.4 – 1.49 (1.52 – 1.49) | 50.0 – 1.75 (1.79 – 1.75) | 43.2 – 1.40 (1.48-4.40) |
| **Space Group** | I422 | I222 | $P2_12_12_1$ |
| **Unit cell (a, b, c / Å)** | 90.75, 90.75, 51.94 | 63.02, 88.10, 108.89 | 47.64, 57.74, 65.44 |
| **α, β, γ (°)** | 90, 90, 90 | 90, 90, 90 | 90, 90, 90 |
| **Total reflections** | 442378 (22642) | 391404 (20040) | 421665 (63963) |
| **Unique reflections** | 17998 (998) | 30950 (1499) | 62886 (10033) |
| **Completeness (%)** | 99.8 (99.7) | 99.9 (99.8) | 99.8 (98.7) |
| **$I / \sigma I$** | 25 (0.5) | 39.2 (1.5) | 10.1 (0.9) |
| **$R_{merge}$** | 0.052 (1.55) | 0.026 (1.57) | 0.104 (1.18) |
| **R-meas** | 0.055 (1.70) | 0.029 (1.88) | 0.112 (1.60) |
| **Multiplicity** | 24.6 (21.5) | 12.9 (13.4) | 14.9 (15.0) |
| **Reflections in refinement** | 17998 | 30839 | 33210 |
| **Reflections used for R-free** | 889 | 1504 | 1646 |
| **R-work** | 0.254 | 0.251 | 0.173 |
| **R-free** | 0.270 | 0.315 | 0.207 |
| **RMS bonds** | 0.009 | 0.009 | 0.010 |
| **RMD angles** | 1.70 | 1.67 | 1.60 |
| **Ramachandran favored (%)** | 96 | 94 | 98 |
| **Ramachandran allowed (%)** | 4 | 4 | 2 |
| **Ramachandran outliers (%)** | 0 | 2 | 0 |

**Supplementary Table 2. Crystal data collection and refinement statistics**
Structures were collected and solved from a single crystal, values in parentheses are for highest-resolution shell.

**Renko et al, Supplementary Table 2**
